# Applying genomic data to seagrass conservation

**DOI:** 10.1101/2020.08.18.255307

**Authors:** Nikki L Phair, Erica S Nielsen, Sophie von der Heyden

**Affiliations:** Evolutionary Genomics Group, Department of Botany and Zoology, University of Stellenbosch, Private Bag X1, Matieland, 7602, South Africa

## Abstract

Although genomic diversity is increasingly recognised as a key component of biodiversity, it is seldom used to inform conservation planning. Estuaries and keystone species such as the southern African seagrass, *Zostera capensis*, are under severe anthropogenic pressure and are often poorly protected. In this study we integrated SNP data generated from populations of *Z. capensis* across the South African coastline into the spatial prioritisation tool Marxan. We included different measures of genomic variation to account for genomic diversity, distinctness and evolutionary potential to explore spatial planning scenarios. We investigated how conservation priority areas identified by targeting only habitat type, differed from those identified by also including genomic measures; further we assessed how different genetic diversity metrics change prioritisation outcomes. All scenarios targeting genomic variation identified unique conservation prioritisation areas compared to scenarios only targeting habitat type. As such, omitting these estuaries from regional MPA networks risks the loss of evolutionarily important populations, threatening resilience and persistence of associated estuarine communities and their ecosystem services. We also observed a high degree of overlap between prioritisation outcomes across targeted measures of genomic variation. As such, by including even single measures of genomic variation, it may be possible to sufficiently represent the evolutionary processes behind the patterns of variation, while simplifying the conservation prioritisation procedure.

## Introduction

Estuaries are some of the most dynamic coastal systems globally and are important for supporting high levels of biodiversity and foundational species such as saltmarshes, mangroves and seagrass meadows, which provide vital ecosystem services including elevated levels of carbon sequestration, supporting regional fisheries and maintaining water quality (Bertelli and Unsworth 2014; Jackson et al. 2015). However, most coastal ecosystems, including estuaries, are under intense anthropogenic pressures due to their proximity to human populations (Mead et al. 2013; Little et al. 2017). Further, these systems are expected to be at increased risk of habitat degradation in future decades due to climate change, which will only be further exacerbated by human induced pressures. South Africa is no exception, even accounting for the progress made in establishing 20 new MPAs, for a total of 42 (Skowno et al. 2019). Yet, many of South Africa’s existing coastal MPAs are ineffective, with fishing effort in estuaries up to five times higher inside compared to outside restricted areas (Skowno et al. 2019). This leaves foundational species, such as the seagrass *Zostera capensis*, and their vital ecosystems services vulnerable.

Increased pressures on coastal systems are especially concerning as models project around 30% of seagrass-suitable habitat will be lost along the South African coastline by the year 2070 (Phair 2016). In reality, *Z. capensis* declines may be even more extreme as the effects of climate change are compounded by human pressures (Adams 2016). Indeed, declines and local extirpations of *Z. capensis* have already been recorded (Human et al. 2016) and given its disjunct distribution, once *Z. capensis* is lost in a particular estuary, it is unlikely to be recolonised from elsewhere (Adams 2016). In addition, *Z. capensis* is highly clonal, with distinct evolutionary dynamics that separate two historically divergent clusters (Phair et al. 2019), where lowered genomic diversity is associated with anthropogenic pressures (Phair et al. 2020). The combination of these factors question the persistence of *Z. capensis* in South Africa and are likely to increase the loss of meadows further. In order to safeguard the future of *Z. capensis* and its associated biodiversity and human benefits, increasing the number and quality of marine and coastal protected habitat is crucial.

The extent to which protected areas succeed in protecting biodiversity relies primarily on meeting two key objectives, representativeness and persistence (Margules & Pressey 2000). Protected areas should capture a representative sample of the full range of biodiversity across levels of organisation including genetic diversity, which is increasingly recognised as a key component of biological diversity. In fact, evidence is mounting that including genetic data into conservation planning is essential to incorporate evolutionary potential and enhance contemporary and future management of natural resources (Beger et al. 2014; Carvalho et al. 2017; Nielsen et al. 2017; Diniz-Filho et al. 2018; Paz-Vinas et al. 2018). Evolutionary potential is an important facet in conservation planning, as it underpins the capacity of species and populations to adapt to and persist through changing conditions (Mittell et al. 2015; Rey et al. 2016; Paz-Vinas et al. 2018). For example, Carvalho et al. (2017) demonstrated an increase in biological representativeness and persistence when including measures of evolutionary potential in conservation planning of amphibian and reptile species.

Genomic techniques, specifically those based around Single Nucleotide Polymorphism (SNP) identification have led to significant advances in marine conservation as they can better account for resilience and persistence (compared to for example mitochondrial DNA or microsatellites, that predominantly provide insights into neutral evolutionary processes), and improve our understanding of the mechanisms behind adaptation and speciation (Nielsen et al. 2009; Allendorf et al. 2010; Reitzel et al. 2013; McMahon et al. 2014; Gaither et al. 2018; Nielsen et al. 2018), as well as connectivity (Nielsen et al. 2017; Rodriguez-Ezpeleta et al. 2017). Both standing genomic diversity and local adaptation can increase evolutionary resilience, and should therefore be considered when planning MPAs with the aim of enhancing species persistence through climate change (Sgrò et al. 2011; Bible & Sanford 2016). For this purpose, it may be useful to apply a genomic approach to the conservation of a keystone estuarine species such as *Z. capensis*, which also functions as an umbrella species whose conservation ensures the protection of many other species.

In this study, we utilised an existing SNP data set to explore conservation outcomes for the endangered seagrass, *Zostera capensis* throughout its distribution in South Africa. Specifically, we integrated different measures of genomic variation to account for genomic diversity, distinctness and evolutionary potential using the spatial prioritisation tool Marxan, to assess conservation prioritisation under different genomic scenarios. We looked at 1) how conservation priority areas identified by targeting only habitat differed from those identified by also targeting different genomic measures, and 2) how different genomic metrics change prioritisation outcomes. This study contributes towards building a framework to better understand the impact of different types of genomic data on spatial planning for vulnerable coastal ecosystems.

## Methods

### Data utilised in analyses

Genomic data for this study was generated through a pooled restriction site-associated sequencing (RADseq) approach by Phair et al. (2019) (NCBI PRJNA503110; GeOMe project: *Zostera capensis* pooled RADseq; GitHub: https://github.com/vonderHeydenLab/Zostera-capensis-genomics). In short, *Z. capensis* samples from eight locations (Olifants, Berg, Breede, Knysna, Swartkops, Nahoon, Mngazana and Richard’s Bay Estuary) across the South African distribution of this species was used to generate a pooled sequencing library. A total of 30 leaf samples per location were collected, with sampling spread throughout meadows to minimise accidental sampling of clonal plants (permit number 0028-AAA008-00159; see Rellstab et al. (2017), Christe et al. (2017) and Jahnke et al. (2019) for sampling clonal and partially clonal plants for pool-seq). Post DNA extraction, DNA was equimolarly pooled by location for ezRAD library preparation and sequencing (Toonen et al. 2013; Knapp et al. 2016; Nielsen et al. 2018); sequence alignment and bioinformatics are described in Phair et al. (2019).

### Genomic measures included in spatial prioritisation scenarios

In order to identify conservation priority areas under various scenarios, the following measures of genomic diversity were included in the analysis: nucleotide diversity (N), expected heterozygosity (H), allelic richness (AR), the number of shared SNPs (S) and private SNPs (PS) as a proportion of the number of total SNPs per population. We also included a measure of evolutionary potential in the form of statistical outliers, that may signal site-specific selection. Combined, the metrics cover distinctness and diversity (Hoban et al. 2020), with each able to address unique conservation objectives (Table 1).

**Table 1.**
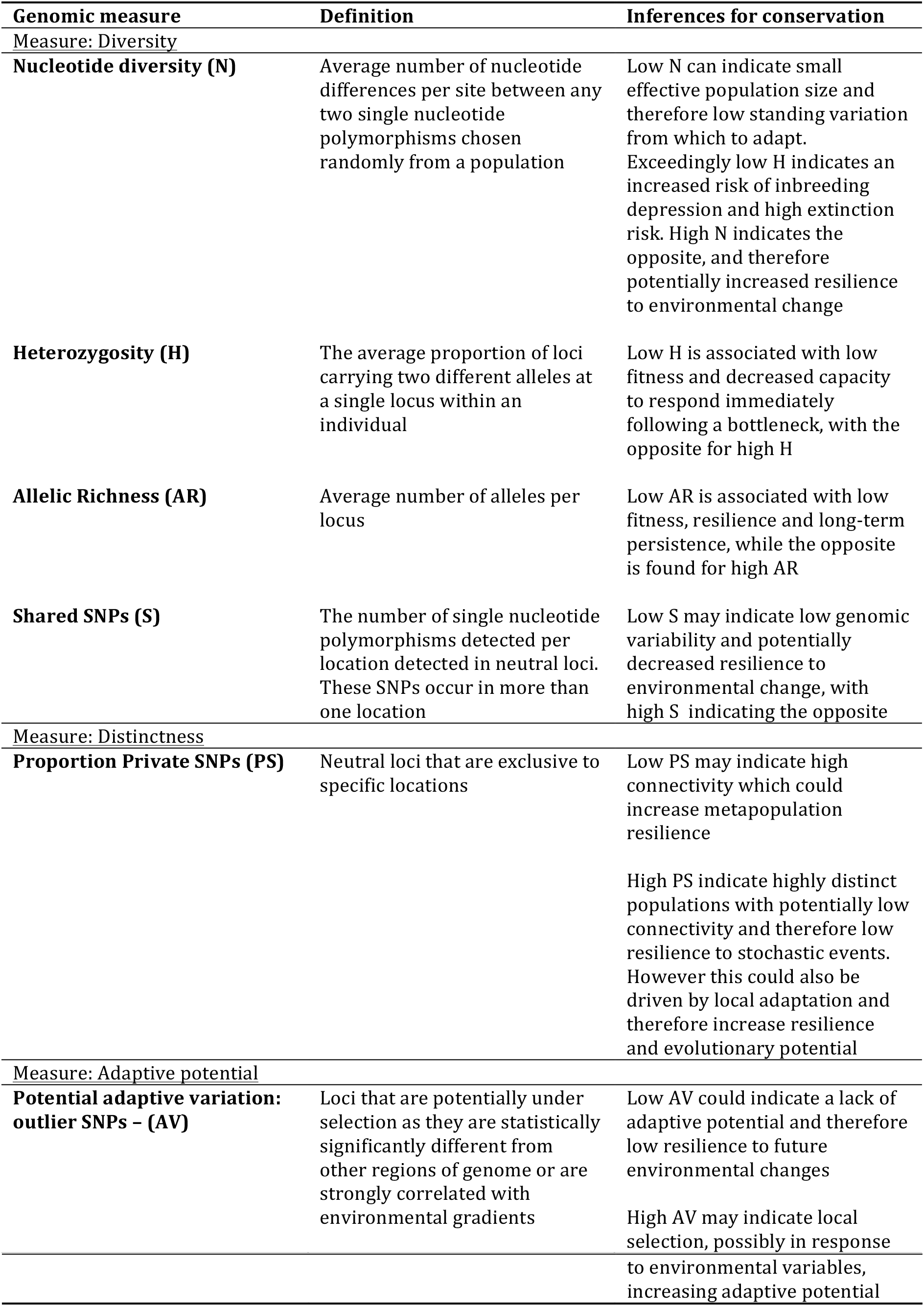
A description of genomic measures included in this analysis and their relevance to conservation prioritisation (after Beger et al. 2014; Nielsen et al. 2017).

### Spatial conservation prioritisation

The decision support tool, Marxan v 2.43 (Ball et al. 2009), was used to design networks of MPAs as possible scenarios for the preservation of *Z. capensis*. Marxan uses an algorithm which minimises reserve cost and size whilst meeting a set of predefined biodiversity targets. South African estuaries were used as planning units. Reserve cost was derived from the fishing effort per estuary quantified in the National Biodiversity Assessment (NBA: van Niekerk et al. 2012) and represents the cost as lost opportunity for industry if MPAs are established as no-take reserves. Habitat data classifying estuaries as permanently open, temporarily open/closed, an estuarine bay or an estuarine lake was obtained from Adams (2016). A baseline scenario was established by targeting 20% of each estuarine habitat type as suggested by the NBA (Skowno et al. 2019), whilst applying the cost layer. Each genomic scenario was developed using the baseline scenario as a foundation (Table 2). The procedures outlined in “Marxan good practices handbook” (Ardron et al. 2010), as well as the methods for integrating genetic data into spatial conservation prioritisation described in Beger et al. (2014), were used to guide the analyses detailed below.

**Table 2.**
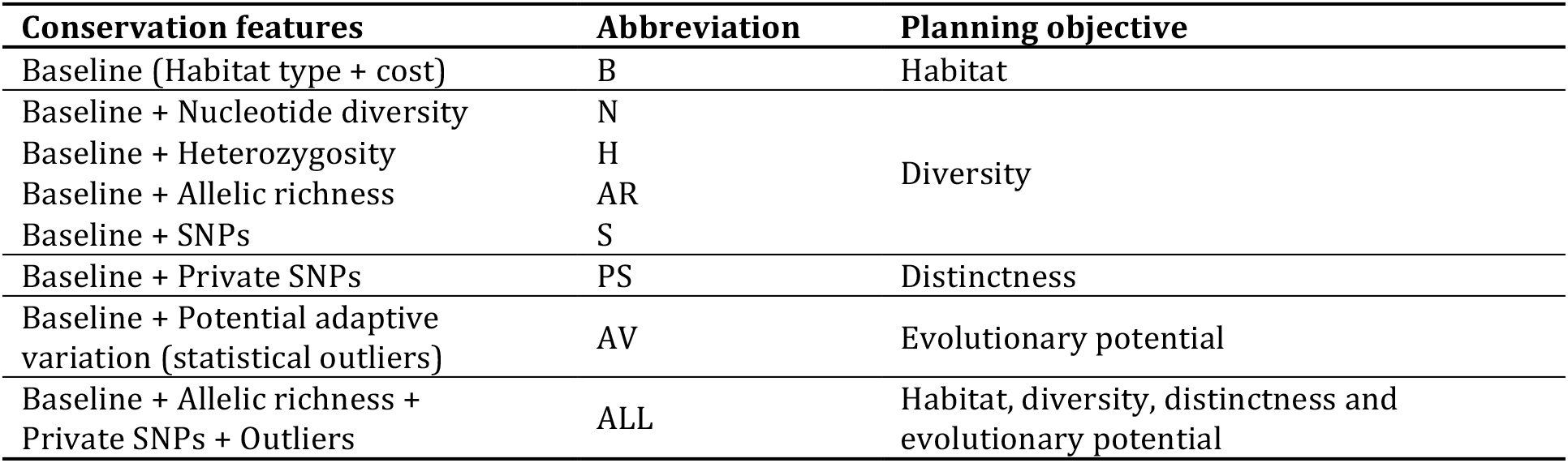
Conservation prioritisation scenarios and planning objectives.

Conservation decision tools such as Marxan require genomic point data to be interpolated throughout the entire planning region to form a spatially continuous surface layer. Accordingly and following Beger et al. (2014) and Nielsen et al. (2017), an Inverse Distance Weighting procedure in ArcMap v10.1 (ESRI, Redlands CA) was used to interpolate genomic data across South African estuaries. The reclassification (reclass) tool in ArcMap v10.1 (ESRI, Redlands CA) was used to reclassify the data from each genomic metric into high, medium and low classes using natural breaks in the data. As both high and low values of genomic diversity are significant in terms of evolutionary processes, targets were set to represent 50% of high and low classes, and 30% of the medium class of each genomic metric following a similar protocol to Beger et al. (2014) and Nielsen et al. (2017).

In addition to the baseline scenario, additional scenarios were run to cover different aspects genomic variability, thus allowing for the comparison of the use of different genomic measures in identifying conservation priority areas (Table 2). Combinations of genomic measures were included in order to observe how priority areas identified change with the addition of data. As it is possible for many different configurations of planning units to meet the conservation objectives, each scenario run was repeated 100 times to account for any system variability, allowing Marxan to calculate planning unit selection frequencies and identify the best solution as the one with the lowest cost to target ratio.

QGIS v2.18.4 (“QGIS Development Team. ‘QGIS Geographic Information System. Open Source Geospatial Foundation Project.’” 2012) was used to visualise scenario outcomes by means of the QMarxan plugin v 1.3.1 (Game & Grantham 2008). Planning unit selection frequencies were obtained from the ‘ssol’ (summed solution) outputs and plotted along the South African coastline for each scenario. In order to understand whether different measures of genomic diversity prioritise different regions, unique and shared priority planning units among diversity scenarios (N, H, AR, S) were identified from the ‘best’ solution outputs and plotted.

We then selected one of these measures of diversity, AR, to represent diversity in subsequent analyses alongside a measure of distinctness (PS), putative adaptive variation (AV) and a combination of all (ALL). We chose allelic richness as it is often considered the most useful measure for monitoring even short-term changes in populations, because of its sensitivity to population declines (Schwartz et al. 2007; Hoban et al. 2014; Gedeon et al. 2017; Phair et al. 2020). For this subset of scenarios (AR, PS, AV, ALL), the differences from and similarities to the baseline scenario, in terms of planning units selected, was obtained from ‘best’ solution outputs and plotted to visualise the impact of including measures of genomic diversity, distinctness and potential adaptive variation in conservation planning in addition to habitat data. Planning unit selection frequencies from the ALL scenario were also plotted alongside estuary threat status, as defined by the NBA (van Niekerk et al. 2018), to visualise the overlap between evolutionary potential and estuarine pressures.

In order to view dissimilarities among scenario solutions, we followed the approach in Harris et al. (2014) and applied nonmetric multidimensional scaling (nMDS) ordination based on Jaccard resemblance matrices in the statistical environment R (R Core Development Team 2008) using Rstudio V 0.98.1102 (“RStudio” 2012). Pearson’s correlation tests were carried out on the selection frequency values for each planning unit to quantify spatial similarities between each pair of scenarios.

## Results

### Spatial conservation prioritisation

All scenarios prioritised estuaries for conservation across the coastline, however the baseline scenario selected estuaries at a lower frequencies than scenarios targeting any of the genomic measures (Fig. 1). Although Pearson correlation tests revealed significant (p <0.05) correlations between all scenarios, there were differences in the spatial distribution of prioritised estuaries and the frequency with which they were selected between scenarios targeting genomic measures (Fig. 1). More specifically, H and PS selected fewer estuaries along the west coast, scenarios N and S fewer along the south coast, and scenarios AR, AV and ALL fewer along the east coast, than other scenarios. Only scenario H and ALL selected the prominent St. Lucia estuary on the east coast. Scenarios N, S and SP selected estuaries at a slightly higher frequency than other scenarios (Fig. 1). Hotspots for the conservation of genomic diversity, distinctness and potential adaptive variation exist along the west, south-west and east coasts, as planning units in these regions were selected at high frequencies across genomic scenarios (Fig. 1).

**Figure 1.**
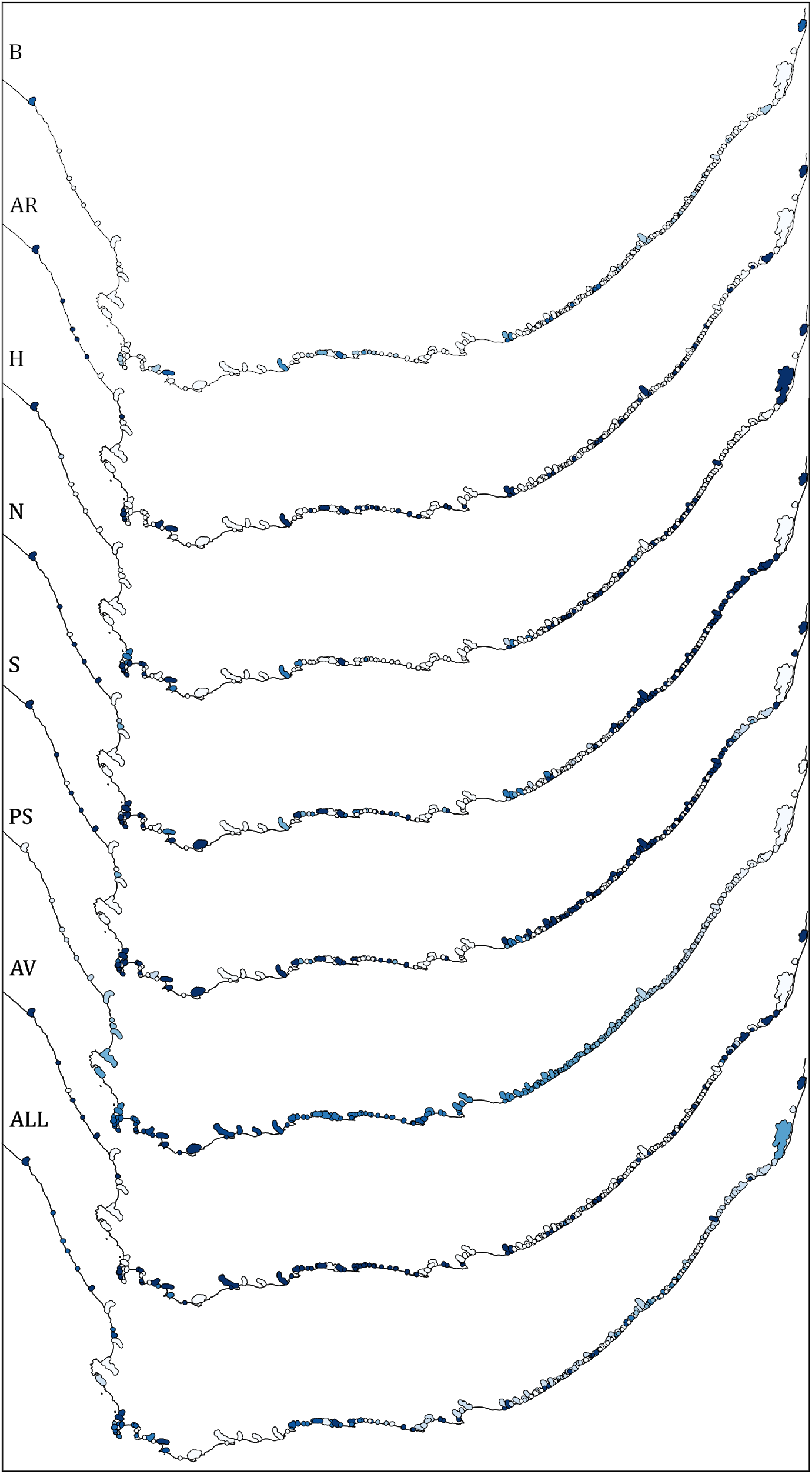
The spatial patterns of selected conservation priority areas across all scenarios with high to low planning unit selection frequency represented by dark to light blue. B = baseline, AR = allelic richness, H = heterozygosity, N = nucleotide diversity, S = SNPs, PS = private SNPs, AV = potential adaptive variation, ALL=combined (also see Table 2).

### Scenario dissimilarities

When scenario dissimilarities were visualised by means of an nMDS plot, the baseline scenario formed a discrete cluster, distant to all other scenarios, with the exception of one anomalous solution (Fig. S1). Solutions from each genomic scenario formed distinct clusters, with solutions from AR and H scenarios most removed from the other genomic scenario clusters, and the ALL scenario displaying the broadest range of solutions. Further, solutions from N, S and PS group closely together, and those from AV fall almost within the ALL cluster of solutions. For many scenarios, such as B and AR, only a few solutions are visible due to highly overlapping nature of these solutions, where Marxan identified the same configuration of priority estuaries in multiple runs.

### Genomic diversity scenarios

Although most prioritised estuaries overlapped across genomic diversity scenarios (AR, H, N and S), each diversity scenario also highlighted unique estuaries for conservation compared to the other diversity scenarios (Fig. 2). Scenario S identified the highest number of unique estuaries for conservation, which were all situated on the south-east coast (Fig. 2). Scenario H only identified one unique estuary for conservation, St. Lucia (Fig. 2). Scenario AR selected unique estuaries for conservation along the south coast and scenario N along the south and east coasts (Fig. 2).

**Figure 2.**
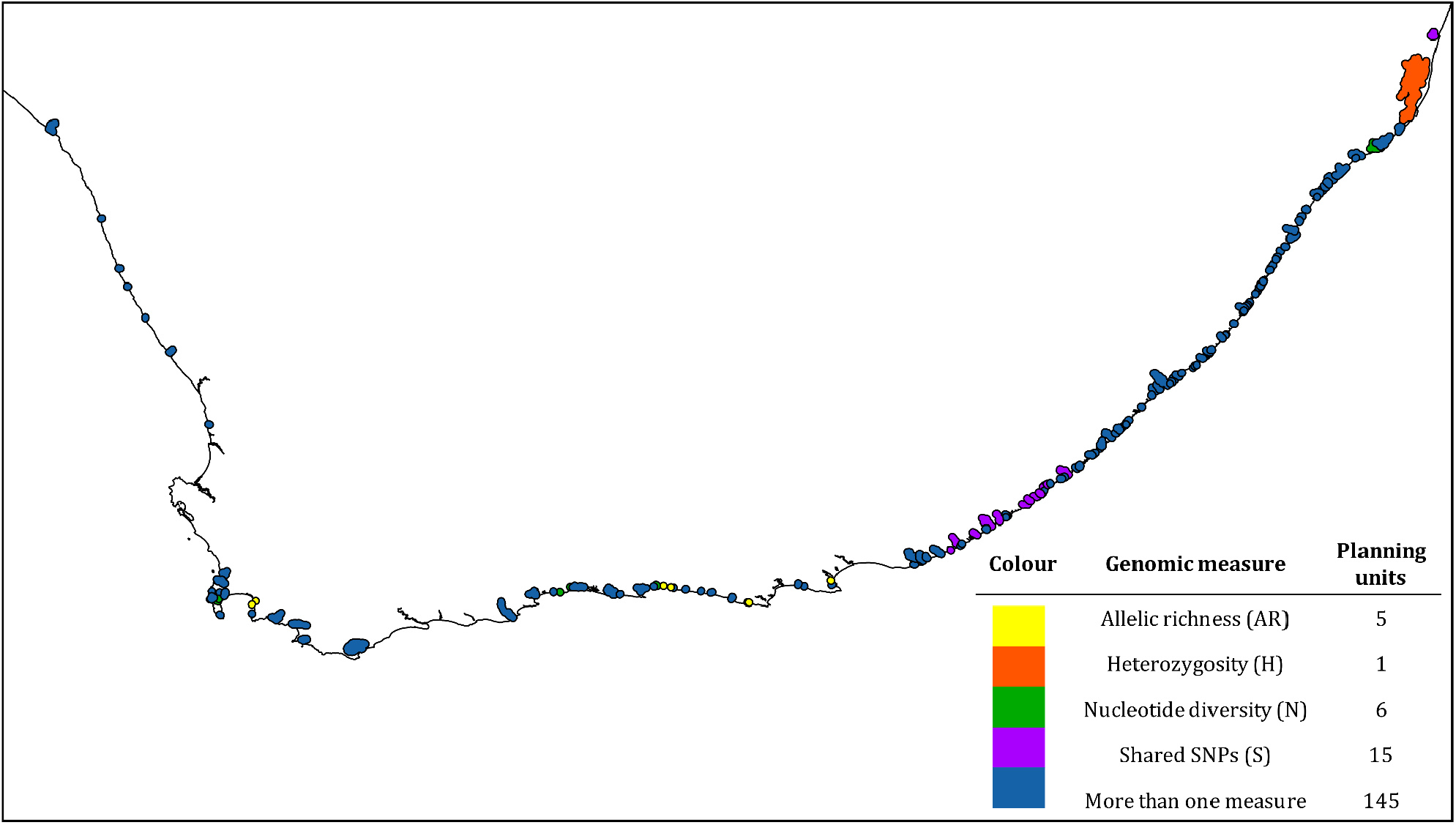
Spatial patterns of selected priority conservation areas derived from conserving habitat as well as diversity measures, with units selected by more than one scenario in blue and those selected only by the scenario based on AR (allelic richness) in yellow, H (expected heterozygosity) in orange, N (nucleotide diversity) in green and S (single nucleotide polymorphisms) in purple.

### Genomic diversity vs distinctness vs potential adaptive variation

The majority of prioritised estuaries were identified across the baseline (habitat) scenario and scenarios targeting diversity (AR), distinctness (PS), potential adaptive variation (AV) and ALL. However, each of these scenarios also identified unique estuaries that were gained or lost compared to the baseline (Figure 4). Scenario PS was the most dissimilar from the baseline scenario, as it showed the greatest number of estuaries gained and lost with respect to those selected by the baseline scenario, which is also evident from the nMDS plot (Fig. S1).

### Overlapping threat status and genomic conservation planning

When comparing the planning unit selection frequency of the ALL scenario with the threat status of estuarine systems, the majority of the critically endangered and vulnerable estuaries are captured (Fig. 4). However, there is little overlap with the current coastal MPA network (Skowno et al. 2019), particularly along the west and south coasts, where many estuaries are classified as critically endangered or endangered, and are highlighted by their evolutionary importance (Fig. 4).

## Discussion

With the threats to coastal systems escalating due to climate change and increasing anthropogenic pressures, MPA networks are vital for the persistence of coastal species and associated ecosystem services. Seagrasses provide a broad array of services that support both biodiversity and human well-being and are thus crucial for coastal ecosystem health, with *Z. capensis* identified as an important keystone species in South African estuarine systems (Adams 2016). Although it is recognised that genomic diversity is important for species and ecosystem resilience (Benestan et al. 2016; Evans et al. 2017a, b; Timpane-Padgham et al. 2017), there are limited examples of evolutionary patterns, particularly potential adaptive variation (Pearse 2016), integrated into actionable conservation and management plans (Sork et al. 2009; Laikre et al. 2009; Laikre 2010; Beger et al. 2014; von der Heyden et al. 2014; Nielsen et al. 2017; von der Heyden 2017; Hoban et al. 2020). One reason for this is because there is no clear evidence for how different genetic and genomic measures vary within a conservation planning framework, hindering their uptake into a more formalised process identifying priority areas. This study provides unique spatial plans that not only compare different metrics that capture genomic diversity, distinctness, and potential signals of local adaptation, but also provide some insights into the feasibility of including these measures into a conservation plan for *Z. capensis* in South Africa. Notably, this study illustrates the importance of including genomic information in MPA planning, and the risk to the evolutionary processes which drive genomic patterns if management plans are based solely on habitat level data.

### Importance of genomic data in spatial planning

Although priority areas overlapped across all scenarios, as they were all founded on baseline habitat data, baseline and genomic scenarios identified noticeably different estuaries for conservation (Fig. 1). This is consistent with findings from other single species (Beger et al. 2014) and even multi-species approaches (Nielsen et al. 2017), and highlights potentially significant omissions in traditional habitat-based MPA design. As genomic diversity is foundational for adaptation and resilience to environmental change, excluding such data from planning seriously undermines future persistence of natural populations (von der Heyden 2009; Beger et al. 2014; von der Heyden et al. 2014). Within the context of our study, this has important implications for future-proofing *Z. capensis* populations along the South African coast, where many estuaries are under intense anthropogenic pressures (van Niekerk et al. 2018; Skowno et al. 2019; Fig. 4). It is essential that genomic data is included as it offers unique insights into the species resilience. Further, failing to conserve current genomic variation of *Z. capensis* increases the probability of losing genotypes which may be more resilient to environmental change. This has already been demonstrated in Phair et al. (2020) where anthropogenic stressors were observed to play an important role in reducing genomic diversity, likely because of the loss of resilient genotypes. This is highly problematic, as numerous seagrass studies suggest that genomic diversity is linked to increased resistance and resilience in various forms (Massa et al. 2013; Jahnke et al. 2015; Phair et al. 2020; Table 1). Although single species approaches in conservation management are often criticised as any one species may not be representative of the broader ecosystem (Richardson et al. 2016; Anthonysamy et al. 2018), their use is recognisably justified when dealing with keystone species such as seagrasses, where conserving such ‘umbrella’ species can ensure the protection of a wide range associated biodiversity (Simberloff 1998; Hughes et al. 2009).

### Measures of genomic diversity, distinctness and adaptation resolve different conservation priorities

Genomic diversity scenarios (AR, H, N, S) all identified statistically similar areas for conservation prioritisation. Between one and 15 priority planning units unique to each scenario were identified, whilst all others were shared among scenarios (Fig. 2). This suggests that conserving a proportion of estuaries with low, medium and high variation for any single genomic diversity measure may sufficiently capture priority estuaries identified by other measures. This has also been observed for measures of genetic diversity (from mtDNA), where Nielsen et al. (2017) consistently identified congruent patterns of spatial prioritisation when targeting haplotype diversity, nucleotide diversity, local genetic differentiation and private alleles, providing support that targeting multiple classes of any one measure of genetic variation can adequately represent the evolutionary patterns observed in other genetic metrics. In this regard, the process of integrating genomic information into spatial planning may be somewhat simplified for conservation managers by employing the most easily obtainable genomic measures.

In addition to measures of diversity it is also important to consider distinctness, where conservation objectives can focus on uniqueness or similarity. For example, highly structured populations may harbour unique genetic variants that could be of conservation concern, whereas less structured populations are likely to have some connectivity and thus be more resilient to perturbations (Chust et al. 2013; Grech et al. 2018). Genomic distinctness scenarios (here measured as private SNPs unique to populations) identified estuaries along the entire coastline. As the scenario targeted various levels of genomic distinctness, these estuaries represent meadows with both high and low levels of connectivity. For example, estuaries on the west coast may be more distinct than those clustered on the south and east coasts, which may be more highly connected (Fig. 3). This approach is beneficial as it safeguards evolutionary potential in a two-pronged approach, firstly, by preserving more homogeneous meadows, which may be more resilient through rescue of declining populations by adjacent well-connected populations (McMahon et al. 2017; Grech et al. 2018) and secondly, by preserving locally adapted populations that may be pre-adapted to specific environmental stressors.

**Figure 3.**
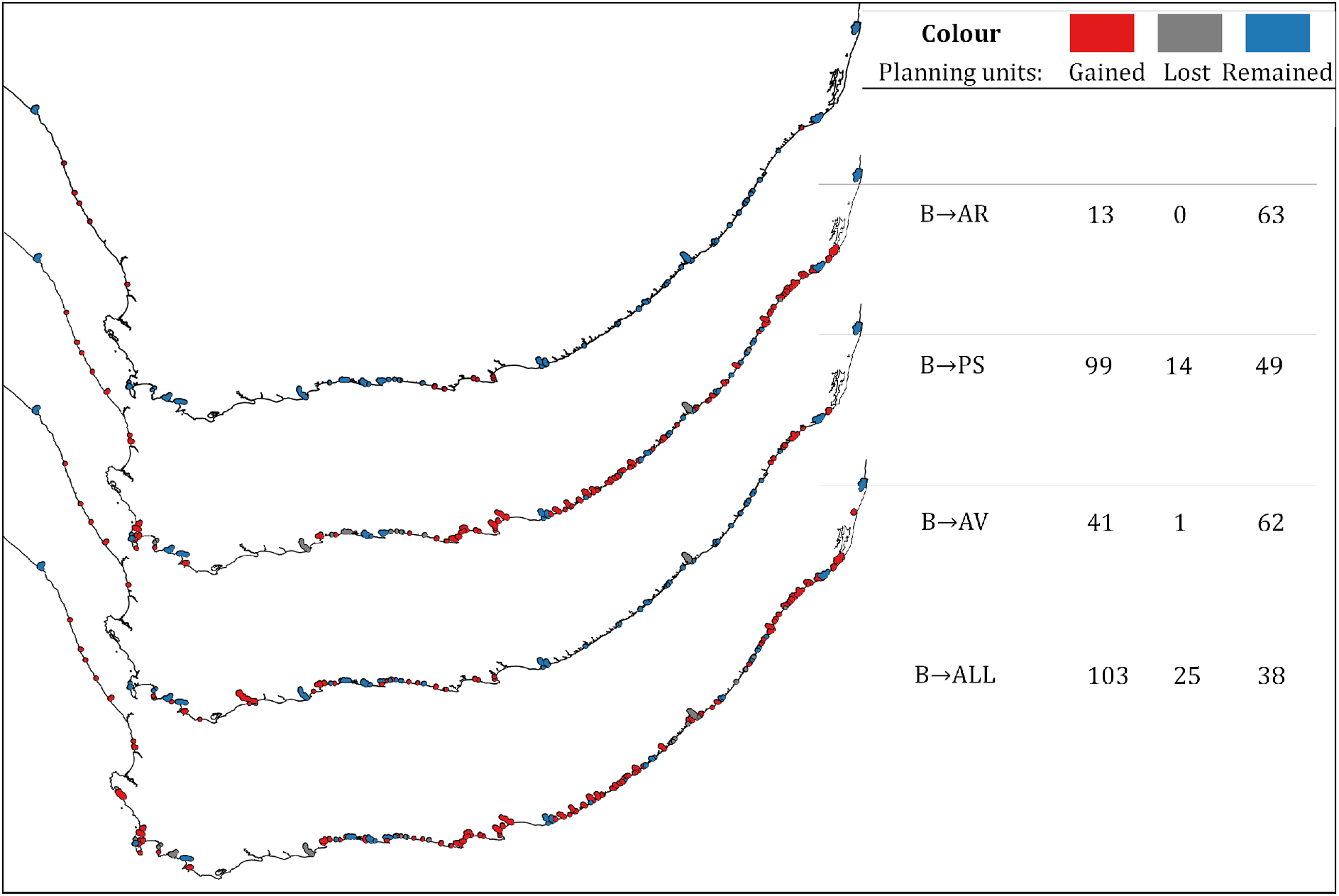
Change in spatial patterns of selected priority conservation areas with the addition of genomic measures of diversity (AR = allelic richness), distinctness (PS = private single nucleotide polymorphisms), potential adaptive variation (AV = adaptive variation), and a combination thereof (ALL) to solely targeting habitat, with units gained in red, lost in grey and remaining selected in blue (number of planning units indicated on the right).

In order to preserve evolutionary potential, it is important to consider potential adaptive variation in addition to distinctness and diversity, as locally adapted genetic variants may exhibit population-specific responses to the environment (Sgrò et al. 2011; Carvalho et al. 2017; Hoban 2018; Razgour et al. 2018). Notably, the scenario targeting potential adaptive variation (AV) identified priority areas distinct from the baseline habitat scenario, on the west, south and north-east coasts (Fig. 3). These regions could represent areas of high adaptive potential and therefore resilience to environmental change, under the assumption that the statistical outliers detected in Phair et al. (2019) are indeed of adaptive importance and have conservation relevance. As different genomic metrics may highlight different priority areas, it is important to carefully choose metrics based on conservation objectives.

Although RAD-seq generated genomic data provides an abundance of evolutionary information, it is also important to acknowledge its limitations, particularly in the potential bias introduced when identifying putative outlier loci (Kofler et al. 2016; Lowry et al. 2017a, b). Even so, the use of genomic data in conservation is increasingly being advocated for (Funk et al. 2012; Hohenlohe et al. 2017; Lowry et al. 2017a), and conservation practitioners should account for such uncertainty in genomic data as they would with other biological data which informs conservation decisions (Kujala et al. 2013).

### *Threats to the evolutionary diversity of* Z. capensis

Our studies shows mismatch between sufficient protection and the distribution and intensity of anthropogenic pressures on estuaries along the coastline (van Niekerk et al. 2018; Fig. 4). The recent expansion of the MPA network is an important step in protecting biodiversity and increasing sustainability, however more focus could still be given to foundational coastal ecosystems, including estuaries. The mismatch between estuarine protection and threat level is particularly evident on the south and south-west coasts, where there are a high number of estuaries rated as endangered and critically endangered in terms of a loss of function and structure due to anthropogenic and climate pressures (Fig. 4). Further, these estuaries correspond to areas identified as priorities for conservation by genomic scenarios (Fig. 4). As such, omitting these estuaries from MPA networks risks the loss of evolutionarily important populations of *Z. capensis* and could threaten the resilience and persistence of not only this keystone species, but also estuarine associated communities. Going forward, MPAs should be reviewed in order to ensure persistence and representativeness of evolutionary potential of keystone species, and thus estuarine associated communities and ecosystem services, in a cost-effective manner.

**Figure 4.**
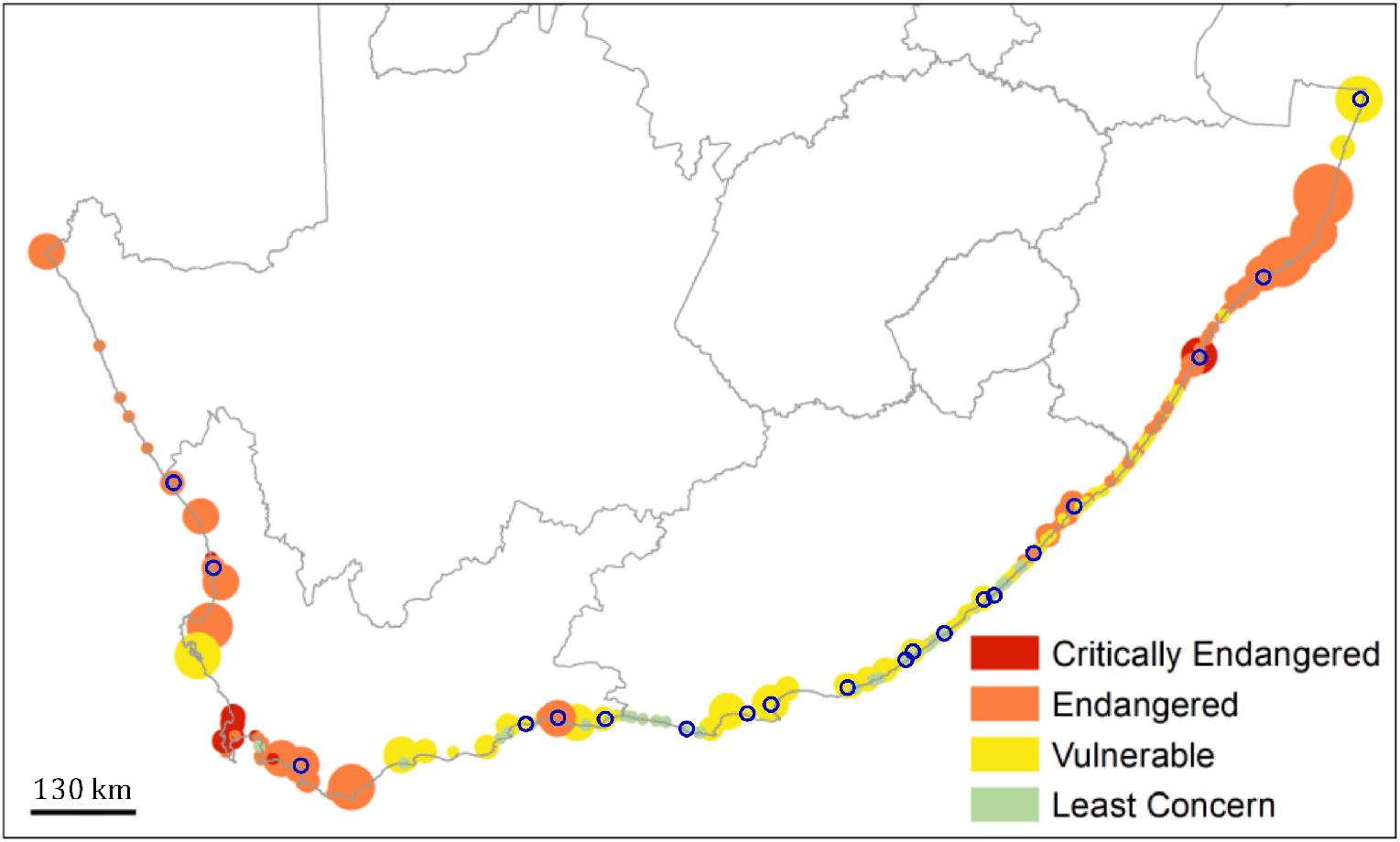
Threat status of South African estuaries (from the NBA; van Niekerk et al. 2018) with the most highly prioritised estuaries under the ALL scenario circled in blue.

