## Supplemental Figure 1 for "Applying genomic data to seagrass conservation"

### Supporting Information

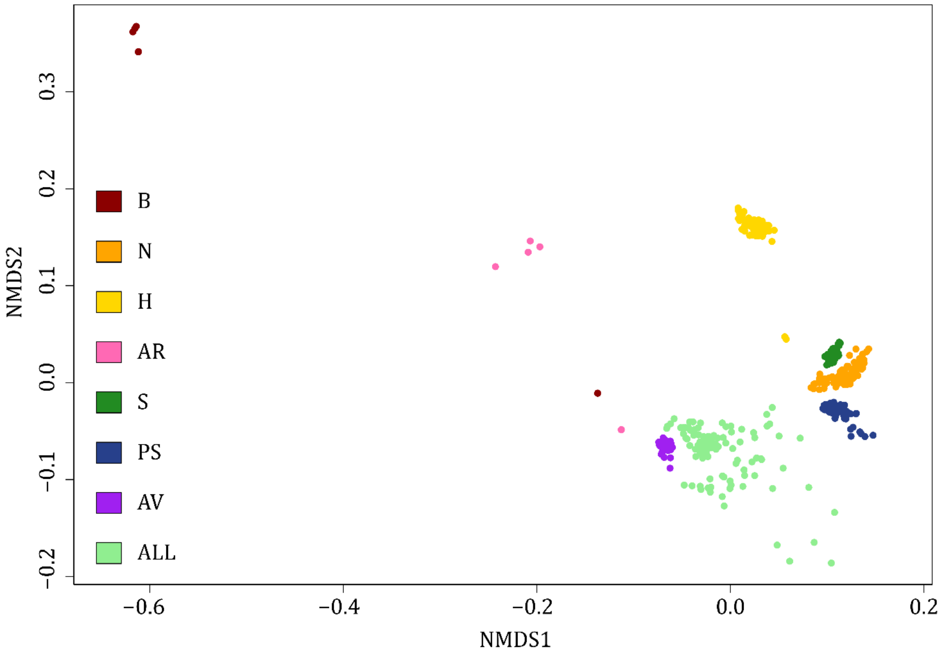


**Figure S1** nonmetric multidimensional scaling plot displaying dissimilarities among scenarios. B = baseline; N = nucleotide diversity; H = expected heterozygosity; AR = allelic richness; S = single nucleotide polymorphisms; PS = private single nucleotide polymorphisms; AV = adaptive variation; ALL = AR, PS and AV combined scenario.
